# A cancer-associated missense mutation in PP2A-Aα increases centrosome clustering during mitosis

**DOI:** 10.1101/570978

**Authors:** Noelle V. Antao, Marina Marcet-Ortega, Paolo Cifani, Alex Kentsis, Emily A. Foley

## Abstract

A single incidence of whole-genome doubling (WGD) is common early in tumorigenesis. In addition to increasing ploidy, WGD doubles centrosome number. In the ensuing mitoses, excess centrosomes form a multipolar spindle, resulting in a lethal multipolar cell division. To survive, cells must cluster centrosomes into two poles to allow a bipolar cell division. Cancer cells are typically more proficient at centrosome clustering than untransformed cells, but the mechanism behind increased clustering ability is not well understood. Heterozygous missense mutations in *PPP2R1A*, which encodes the alpha isoform of the A-subunit of protein phosphatase 2A (PP2A-Aα), positively correlate with WGD. To understand this correlation, we introduced a heterozygous hotspot mutation, P179R, in endogenous PP2A-Aα in human tissue culture cells. We find that PP2A-Aα^P179R^ decreases PP2A assembly and targeting. Strikingly, when centrosome number is increased, either through cytokinesis failure or centrosome amplification, PP2A-Aα mutant cells are more proficient than WT cells at centrosome clustering, likely due to PP2A-Aα loss-of-function. PP2A-Aα^P179R^ appears to enhance centrosome clustering by altering the interactions between centrosomes and the cell cortex. Thus, cancer-associated mutations in PP2A-Aα may increase cellular fitness after WGD by enhancing centrosome clustering.

## Introduction

Most human tumors are aneuploid. In about one-third of tumors, aneuploidy is preceded by a whole-genome doubling (WGD) (Zack et al., 2013), an event that correlates with poor prognosis for patients (Bielski et al., 2018). After WGD, the newly tetraploid genome promotes the evolution of aneuploid karyotypes (Dewhurst et al., 2014), with tumor cells losing chromosomes and acquiring chromosomal rearrangements (Carter et al., 2012; Zack et al., 2013). However, WGD can be detrimental to subsequent mitoses because centrosome number doubles (from 2 to 4) concomitantly with genome doubling. Because each centrosome nucleates microtubules during mitosis (Ring et al., 1982), a multipolar spindle forms, which can result in a lethal multipolar cell division (Ganem et al., 2009). Centrosome amplification is nevertheless common in cancers (Chan, 2011) because cells can cluster supernumerary centrosomes into a pseudo-bipolar spindle (Quintyne et al., 2005). Because lethal multipolar cell divisions are expected to decrease the fitness of a tumor cell population, cancer cells likely experience selective pressures to efficiently cluster supernumerary centrosomes.

Centrosome clustering occurs via motor proteins located near spindle poles and centrosomes, such as dynein and KIFC1/HSET, as well as proteins localized at the kinetochore/centromere that control microtubule binding and spindle assembly checkpoint signaling (Drosopoulos et al., 2014; Kwon et al., 2008; Leber et al., 2010; Quintyne et al., 2005). At the cell cortex, motor proteins associated with the cortical actin network, such as Myo10 and dynein, position centrosomes through contacts with astral microtubules (Kwon et al., 2015; Quintyne et al., 2005). These proteins contribute to spindle assembly and placement in cells with two centrosomes, suggesting that centrosome clustering occurs through normal cellular pathways. However, centrosome clustering is more efficient in cancer cells compared to non-transformed cells (Ganem et al., 2009). Centrosome clustering efficiency can be increased by depletion of the microtubule binding protein NUMA (Quintyne et al., 2005) or the adhesion protein E-cadherin (Rhys et al., 2018). However, it remains largely unclear how cancer cells evolve to cluster centrosomes more proficiently.

Genomic analyses have identified genetic alterations that positively correlate with WGD (Bielski et al., 2018; Zack et al., 2013). Several changes (*TP53* and *RB1* mutation, *CCNE1* amplification) enable cell proliferation after WGD by weakening the G1 arrest triggered following cytokinesis failure (Andreassen et al., 2001). The molecular impact of other genetic changes associated with WGD, including alterations in protein phosphatase 2A (PP2A), is not known. PP2A is a major source of serine/threonine phosphatase activity and is involved in many cellular processes (Wlodarchak and Xing, 2016). PP2A substrate selectivity arises through protein-protein interactions. In the PP2A heterotrimer, a catalytic subunit (PP2A-Cα/β) and a ‘scaffolding’ subunit (PP2A-Aα/β) are targeted to substrates by families of regulatory subunits. PP2A inactivation was first linked to tumorigenesis when it was discovered that SV40 small t antigen blocks the binding of PP2A-Aα/β to regulatory subunits (Pallas et al., 1990), leading to cellular transformation (Chen et al., 2004). Potentially similar perturbations have been found in tumors, including homozygous deletion of *PPP2R2A*, which encodes the B55α regulatory subunit, and heterozygous missense mutations in *PPP2R1A*, which encodes the α isoform PP2A-A that accounts for ∼90% of total PP2A-A (Zhou et al., 2003). Hotspot mutations in PP2A-Aα are expected to prevent B55 and B56 regulatory subunit binding (Cho and Xu, 2007; Xu et al., 2006; Xu et al., 2008) and likely decrease the functionality of PP2A-Aα. While the impact of PP2A-Aα missense mutations after WGD has not been examined, over-expression of hotspot PP2A-Aα mutants in tissue culture cells has been observed to alter phospho-signaling in other cellular contexts, suggesting the mutations may possibly confer new functions (Haesen et al., 2016; Jeong et al., 2016).

Here, we examine two prevalent hotspot mutations in *PPP2R1A*, P179R and R183W. We find that both mutations reduce PP2A protein-protein interactions. When introduced into one allele of endogenous *PPP2R1A*, the P179R mutation reduces PP2A holoenzyme assembly and intracellular targeting. Strikingly, we find that these changes are sufficient to increase centrosome clustering, in part through factors at the cell cortex that position centrosomes.

## Results

### Characterization of prevalent cancer-associated PP2A-Aα missense mutations

*PPP2R1A* is most frequently mutated in uterine carcinosarcomas (Fig. 1A). To explore the cellular impact of the two most frequent *PPP2R1A* missense mutations (Fig. 1B), we generated retinal pigment epithelial (RPE-1) hTERT cell lines expressing GFP-tagged PP2A-Aα wild type (WT), P179R, or R183W mutants. Each construct was expressed at 30-40% of the level of endogenous PP2A-Aα/β (Fig. 1C). Using quantitative mass spectrometry, we compared the composition of PP2A complexes isolated from SILAC labeled cells by immunoprecipitation of WT or mutant GFP-PP2A-Aα. We focused on proteins with ≥ 2-fold change in binding and P < 0.05 (two-sample t-test). The P179R mutation significantly reduced PP2A-Aα binding to four B56 regulatory subunits (B56α/*PPP2R5A*, B56β/*PPP2R5B*, B56δ/*PPP2R5D* and B56ε/*PPP2R5E*). Accordingly, the binding of proteins that associate with PP2A-Aα via B56 subunits including GEF-H1 (*ARHGEF2*), Liprinα1 (*PPFIA1*) (Hertz et al., 2016), and *MTCL1* (Hyodo et al., 2016) was similarly reduced (Fig. 1D). The P179R mutation also significantly reduced binding to the *B55δ/PPP2R2D* regulatory subunit (Fig. 1D). The binding of STRN regulatory subunits (*STRN, STRN3*, and *STRN4*), a B’’’ regulatory subunit (*PPP2R3A*), and PP2A-C (*PPP2CA, PPP2CB*) was unaffected (Fig. 1D). The R183W mutant had an overall similar decrease in protein-protein interactions (Fig. 1E). While several proteins were observed to have increased binding to a PP2A-Aα mutant, none were shared between the two mutants (Fig. 1F). Thus, the major impact of both the P179R and R183W mutations is to reduce protein-protein interactions.

**Figure 1.**
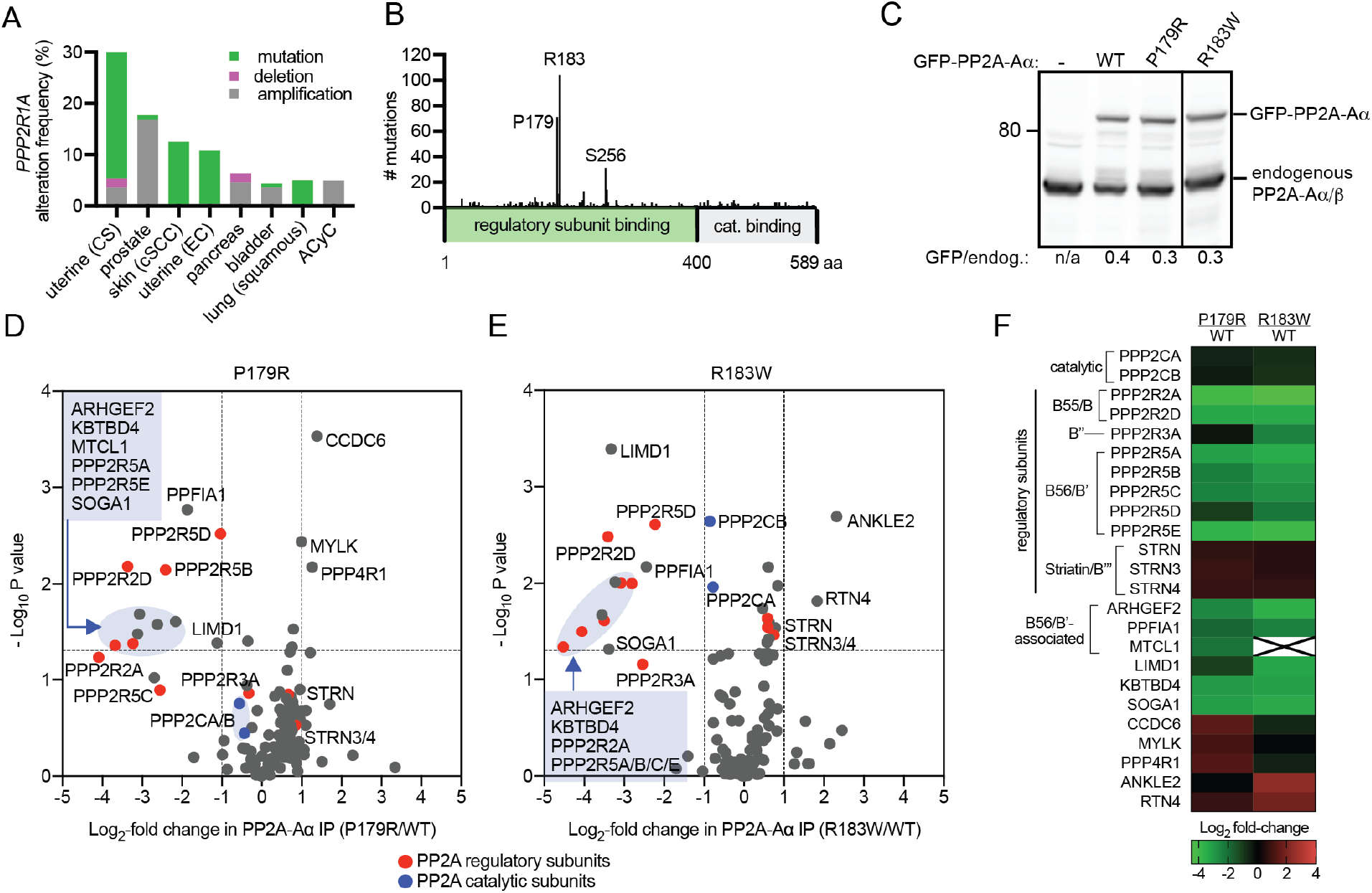
Quantitative proteomic characterization of cancer-associated PP2A-Aα mutations. Incidence of **(A)** *PPP2R1A* alterations **(B)** missense mutations (Cerami et al., 2012). CS, Carcinosarcoma; cSCC, Cutaneous Squamous Cell Carcinoma; EC, Endometrial Carcinoma; ACyC, Adenoid Cystic Carcinoma. **(C)** Western blot analysis of cells expressing GFP-PP2Aα WT or indicated mutants. Solid line indicates intervening lanes have been removed. **(D-E)** Volcano plots with the mean log_2_ fold-change of proteins bound to mutant versus WT PP2Aα against −log_10_ P value. 2-fold change (vertical dashed lines); P <0.05 (horizontal dashed lines); Red and blue circles indicate regulatory and catalytic subunits respectively. **(F)** Heat map of proteins with significant changes in association. Green to red gradient represents the mean log_2_ fold-change. X, protein not detected.

### A heterozygous P179R mutation in PP2A-Aα impacts PP2A holoenzyme assembly in human cells

We next introduced a P179R mutation into one allele of endogenous *PPP2R1A* in RPE-1 cells. This mutation was selected because it is the most common PP2A-Aα missense mutation in uterine tumors, which have the highest incidence of PP2A-Aα alterations (Fig. 1A) (Cerami et al., 2012). We used adeno-associated virus-mediated gene targeting (Berdougo et al., 2009) to introduce a C to G mutation in exon five of *PPP2R1A* (Fig. 2A). We isolated two independent heterozygous clones (Fig. 2B). The mutation did not alter the levels of PP2A-Aα or PP2A-Aα/β (Fig. 2C). Similarly, PP2A-Aα immunoprecipitates from WT and PP2A-Aα^P179R/+^ cells had equivalent levels of PP2A-C (Fig. S1A) and phosphatase activity (Fig. S1B). By contrast, we observed near 2-fold reductions in PP2A-Aα association with B56γ, δ, and ε (Fig. 2D-2E) and B55α (Fig. S1C-D). Consistent with this reduction, mitotic targeting of PP2A^B56^ to the kinetochore/centromere was reduced in PP2A-Aα^P179R/+^ cells (Fig. 2F-G). Collectively, these results indicate that a heterozygous P179R mutation in PP2A-Aα is sufficient to alter the level of a subset of PP2A holoenzymes.

**Figure 2.**
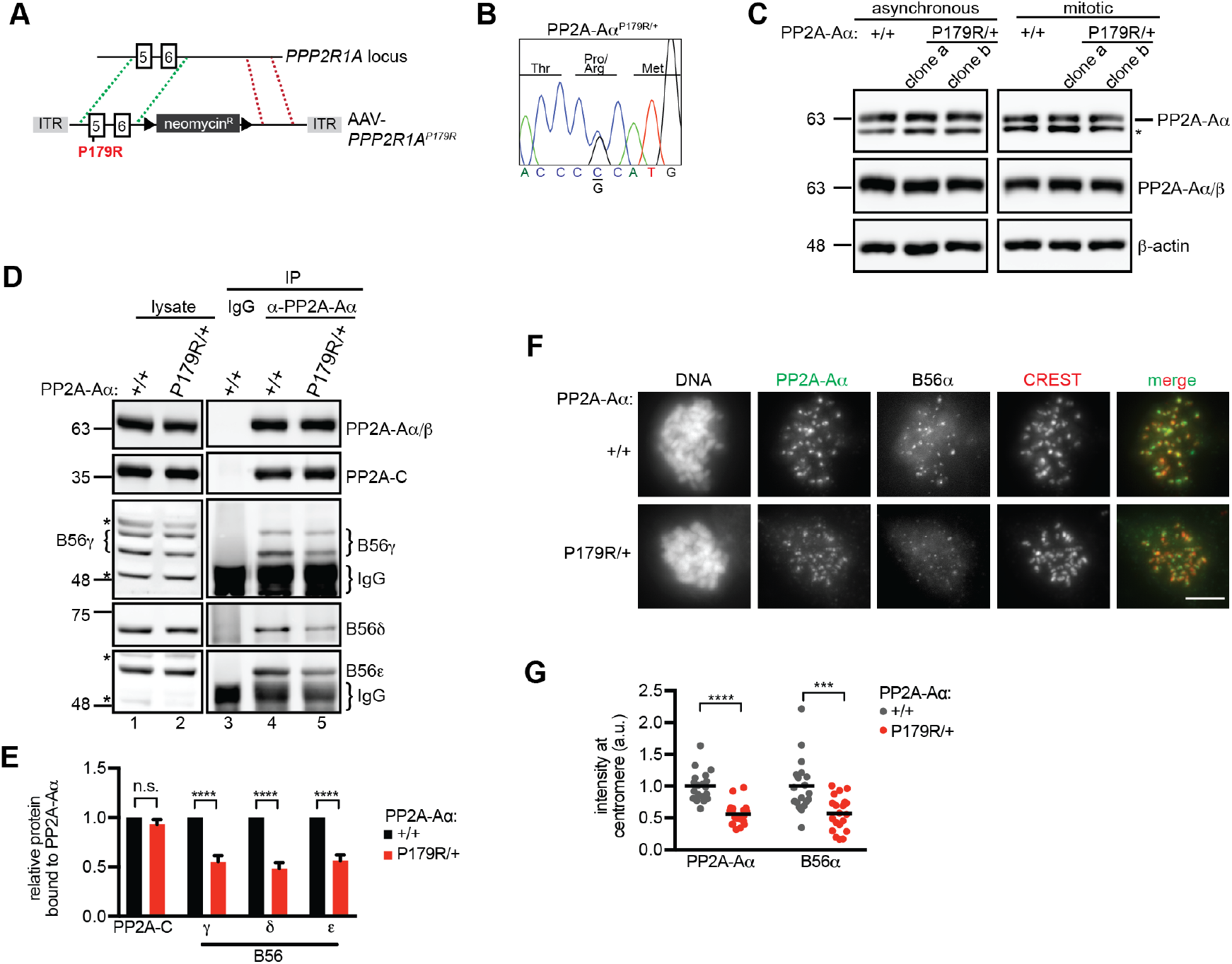
A heterozygous P179R mutation in PP2A-Aα decreases PP2A^B56^ assembly and targeting. **(A)** Schematic of *PPP2R1A* gene-targeting strategy. Exons (rectangles); *loxP* sites (triangles); ITR, AAV-specific Inverted Tandem Repeats; Homologous sequences (green and red dashed lines). **(B)** Sanger sequencing the modified region of *PPP2R1A* in a mutant clone. **(C)** Western blot analysis of lysates from WT (+/+) and independently derived PP2A-Aα^P179R/+^ (P179R/+) clones. *, nonspecific band. **(D)** Western blot analysis of lysates (lanes 1-2) and IPs of control IgG (lane 3) or PP2A-Aα IgG (lane 4-5) in indicated cell lines **(E)** Plotted is the normalized mean + s.e.m. of the experiment in (D) performed three times. **(F-G)** WT (+/+) and PP2A-Aα^P179R/+^ (P179R/+) cells in nocodazole were fixed and processed for immunofluorescence. **(F)** Maximum intensity projection. Scale bar, 5 μm. **(G)** Normalized centromere/kinetochore signal is plotted. Circle, cell; Line, mean. Result is representative of three experiments. a.u., arbitrary units; ns, not significant (P > 0.05),***, P <0.0005,****, P <0.00005 calculated from Student’s two-tailed *t*-test.

### PP2A-Aα^P179R/+^ cells suppress multipolar cell division after cytokinesis failure

To determine if the reduction in PP2A holoenzyme levels might impact PP2A-Aα^P179R/+^ cells after WGD, we induced one round of cytokinesis failure (Yang et al., 2008) by treating cells with cytochalasin D or B, inhibitors of actin polymerization (Cooper, 1987), or with blebbistatin, a myosin II inhibitor (Straight et al., 2003). Binucleate cells were imaged live during the ensuing mitosis (Fig. 3A). In this assay, cytokinesis failure triggers p53 activation causing a G1 cell cycle arrest (Andreassen et al., 2001). Therefore, we treated *Tp53^+/+^* cells with p38 inhibitor SB 203580 (Cuenda et al., 1995) or utilized *Tp53^−/−^* cells. 30% of WT cells treated with cytochalasin D underwent multipolar cell divisions. Strikingly, PP2A-Aα^P179R/+^ cells exhibited 4-fold reduction in the frequency of multipolar cell divisions compared to WT cells (7% in clone a and 8% in clone b) (Fig. 3B-C). PP2A-Aα^P179R/+^ cells also underwent fewer multipolar cell divisions when cytokinesis failure was induced with cytochalasin B or blebbistatin (Fig. 3C). Similar results were observed in *Tp53^−/−^* PP2A-Aα^P179R/+^ cells (Fig. 3D). Expression of GFP-PP2A-Aα-WT, but not the P179R mutant, in PP2A-Aα^P179R/+^ cells partially rescued the cell division phenotype. Expression of PP2A-Aα-P179R in WT cells had no impact on the outcome of cell division (Fig. 3E-F). These data suggest that enhanced bipolar cell division after WGD in PP2A-Aα^P179R/+^ cells is likely due to haploinsufficiency.

**Figure 3.**
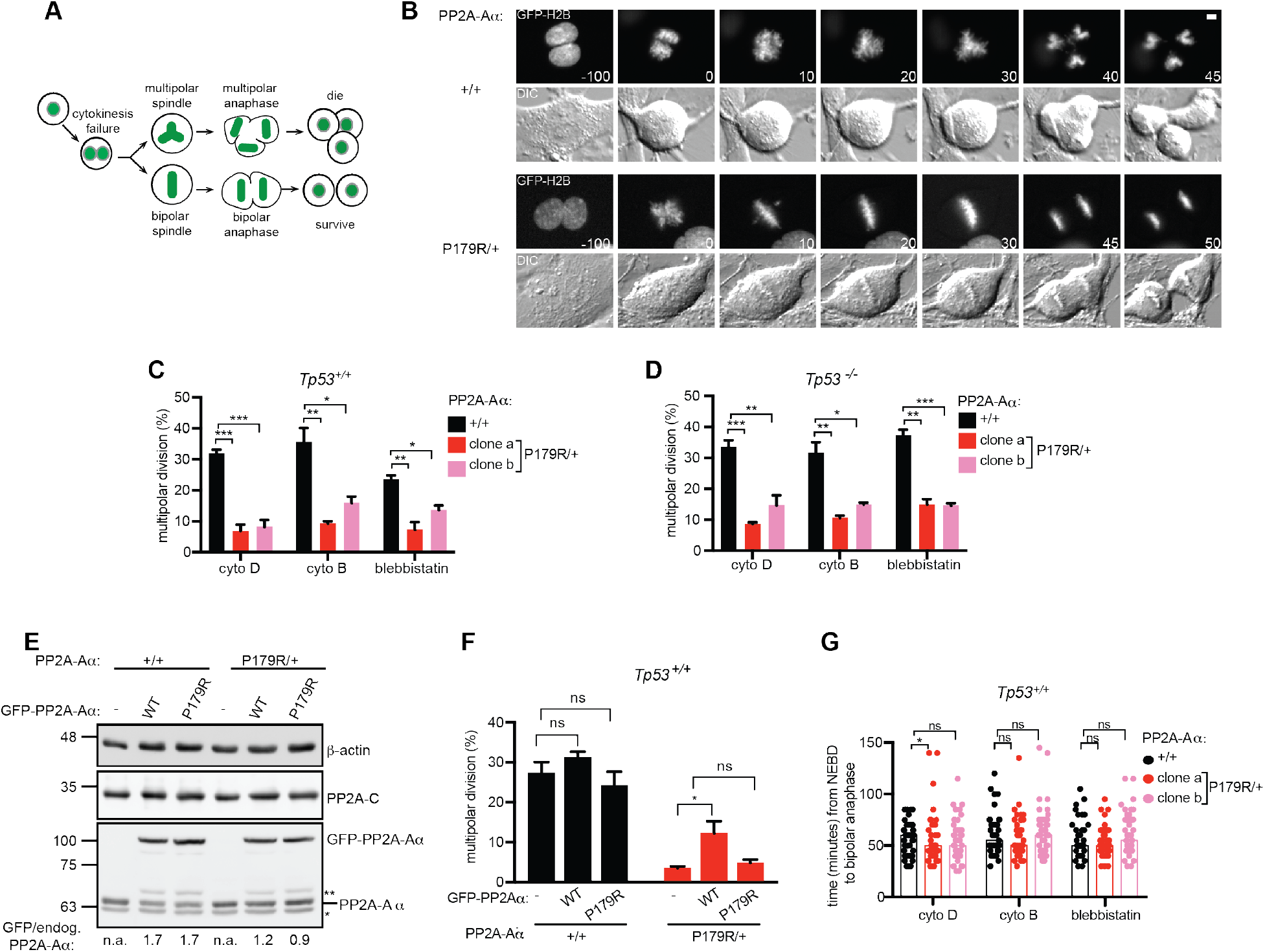
The P179R mutation in PP2A-Aα suppresses multipolar cell division after cytokinesis failure. **(A)** Schematic of cytokinesis failure assay. **(B)** GFP-H2B (top) and differential interference contrast (DIC, bottom) time-lapse images of WT (+/+) and PP2A-Aα^P179R/+^ (P179R/+) cells after cytochalasin D treatment. Time (min) relative to nuclear envelope breakdown is indicated. Scale bar, 5 μm. **(C)** Quantification of cell division outcome of *Tp53^+/+^* cells. **(D)** Quantification of multipolar cell divisions in *Tp53^−/−^* cells treated as in (A). **(E-F)** Cell lines of the indicated genotype expressing GFP-PP2A-Aα-WT or GFP-PP2A-Aα-P179R were **(E)** analyzed by western blotting. *, non-specific band, **, GFP-PP2A-Aα truncation product and **(F)** treated with cytochalasin D as in (A), imaged and cell division outcome quantified. **(G)** Mitotic duration of *Tp53^+/+^* cells in (C) undergoing bipolar cell divisions is plotted. Bars, median; circles, cells. Result is representative of three experiments. (C, D, F) Mean + s.e.m. from three experiments is plotted. * P <0.05, ** P < 0.005, *** P <0.0005, ns, not significant (P >0.05) Student’s *t*-test.

PP2A-Aα^P179R/+^ cells have a modest delay in mitosis (22 min in clone a and 21 min in clone b versus 18 min in WT cells) (Fig. S2A). Because increased time spent in mitosis suppresses multipolar mitoses (Kwon et al., 2008), we compared the mitotic duration of binucleate cells that underwent a bipolar cell division. WT and PP2A-Aα^P179R/+^ cells completed mitosis with indistinguishable kinetics (Fig. 3G) suggesting that the cell division phenotype of PP2A-Aα^P179R/+^ cells is likely due to enhanced bipolar spindle assembly in the presence of extra centrosomes, rather than increased time spent in mitosis.

### PP2A-Aα^P179R/+^ cells cluster supernumerary centrosomes more efficiently

In *Drosophila*, excess centrosomes are sometimes inactivated (Basto et al., 2008). To examine this possibility in PP2A-Aα^P179R/+^ cells, we performed the cytokinesis failure assay and then transiently depolymerized microtubules with nocodazole. After allowing a brief time for microtubule re-growth, we fixed the cells. Microtubules were observed at all centrosomes in all mitotic cells examined (Fig. 4A). Next, we performed the cytokinesis failure assay and identified anaphases with four centrosomes in fixed cells. In PP2A-Aα^P179R/+^ cells, centrosomes were always associated with spindle poles (Fig. 4B) and furthermore, were more likely to be clustered (Fig. 4C).

**Figure 4.**
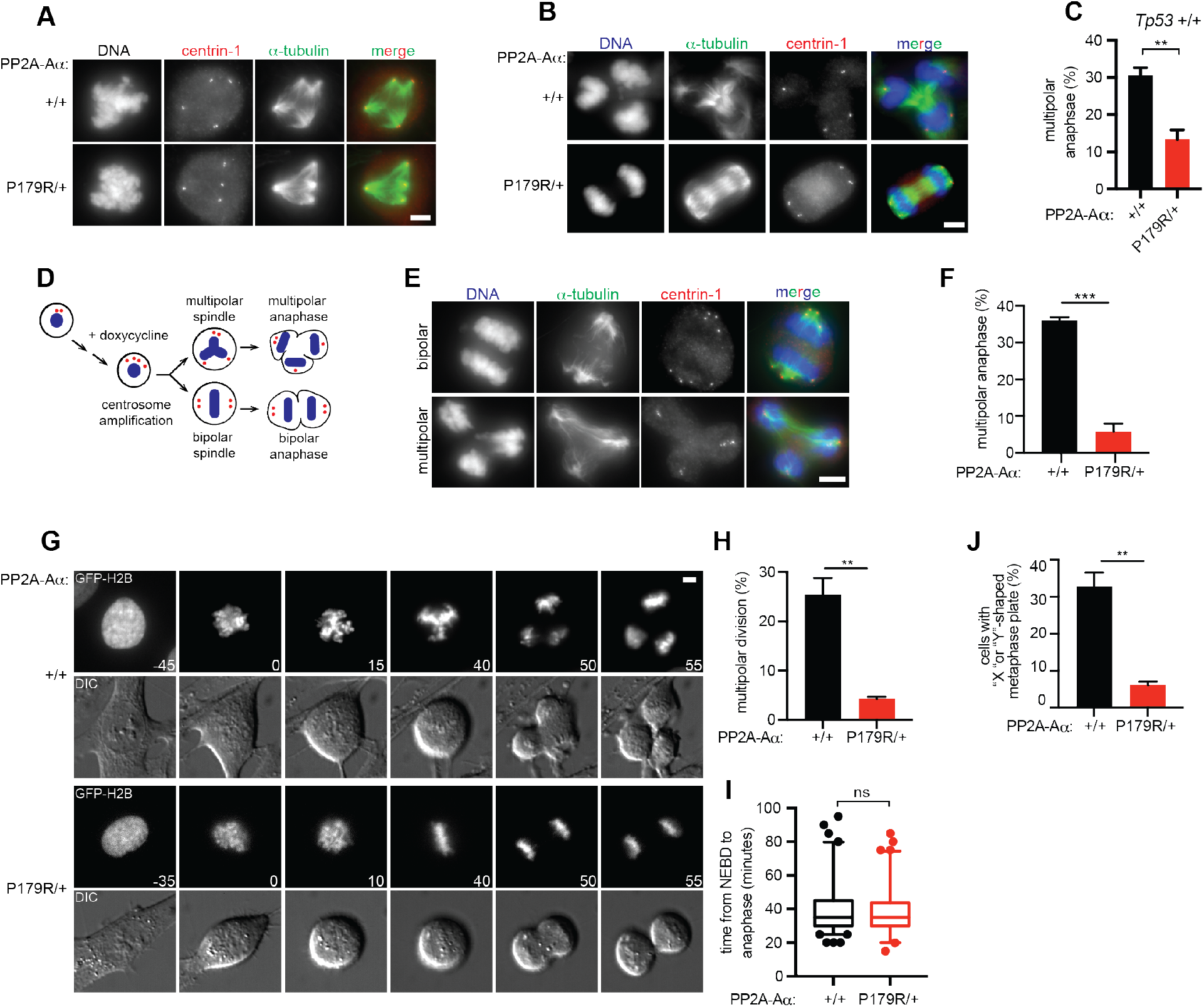
PP2A-Aα^P179R/+^ cells exhibit more robust centrosome clustering. **(A)** Maximum intensity projections of WT (+/+) and PP2A-Aα^P179R/+^ (P179R/+) cells treated with cytochalasin D and then nocodazole. Cells were analyzed by immunofluorescence upon nocodazole removal. **(B-C)** Cells were treated with cytochalasin D and analyzed by immunofluorescence. **(B)** Representative maximum intensity projection. **(C)** Multipolar anaphase incidence in cells with four centrin-1 foci is plotted. **(D)** Schematic of centrosome amplification induction by doxycycline (dox)-inducible Plk4 overexpression. **(E-F)** Plk4-inducible cells were treated with dox and analyzed by immunofluorescence. **(E)** Representative bipolar (top) and multipolar (bottom) anaphases. **(F)** Multipolar anaphase incidence in cells with >4 centrin-1 foci is plotted. **(G-J)** Plk4-inducible cells were treated with dox and imaged live **(G)** GFP-H2B (top) and DIC (bottom) montage. Time (min) relative to nuclear envelope breakdown is indicated. **(H)** Incidence of multipolar cell division is plotted. (I) A box-and-whisker plot of mitotic duration in dox-treated cells undergoing bipolar cell divisions. Circles indicate cells outside of the 5-95 percentile range. Result is representative of three experiments. **(J)** Chromatin configuration was classified as indicative of a multipolar (X or Y shape) or bipolar (I shape) spindle. (C, F, H, J) Mean + s.e.m. from three experiments. Scale bars, 5 μm. **, P <0.005, *** P <0.0005, ns, not significant (P >0.05) Student’s *t*-test.

To test the hypothesis that PP2A-Aα^P179R/+^ cells may cluster supernumerary centrosomes more efficiently than WT cells, centrosome amplification was induced by over-expression of Plk4 kinase (Kleylein-Sohn et al., 2007; Peel et al., 2007) (Fig. 4D). This increases centrosome number without altering ploidy. We introduced doxycycline-inducible Plk4 into *Tp53^−/−^* WT and PP2A-Aα^P179R/+^ cells. After doxycycline induction immunofluorescence analyses of C-Nap1, a marker of functional centrioles (Wang et al., 2011), indicated that 64% of WT cells and 68% of PP2A-Aα^P179R/+^ cells had centrosome amplification (Fig. S2B-C). We then analyzed anaphases of cells with extra centrosomes by immunofluorescence (Fig. 4E). Strikingly, PP2A-Aα^P179R/+^ cells had a 6-fold reduction in multipolar anaphases compared to WT cells (Fig. 4F). A similar result was observed by live imaging (Fig. 4G-H), with no delay in mitosis (Fig. 4I). Additionally, ten minutes prior to anaphase, PP2A-Aα^P179R/+^ cells exhibited a 6-fold reduction in multipolar chromatin configurations (Fig. 4J), suggesting that bipolar spindle geometry is established in advance of anaphase.

### Positioning of centrosomes by cortical forces is enhanced in PP2A-Aα^P179R/+^ cells

To examine how PP2A-Aα^P179R/+^ cells more efficiently cluster centrosomes, we asked if microtubule organization in mitosis is more robust in PP2A-Aα^P179R/+^ cells. Although centrosomes are the major sites of microtubule nucleation, a bipolar spindle can assemble in the absence of one or both centrosomes, albeit with some delay (Khodjakov et al., 2000). We therefore generated acentrosomal cells using centrinone, a small molecule inhibitor of Plk4 (Wong et al., 2015). After centrinone treatment, > 95% of cells had either one or zero centrosomes (Fig. 5A-B). We imaged cells (Fig. 5C) and measured the time between nuclear envelope breakdown and anaphase onset. WT and PP2A-Aα^P179R/+^ cells were similarly delayed upon centrinone treatment (Fig. 5D) indicating that PP2A-Aα^P179R/+^ cells are not more robust at spindle assembly in the absence of centrosomes.

**Figure 5.**
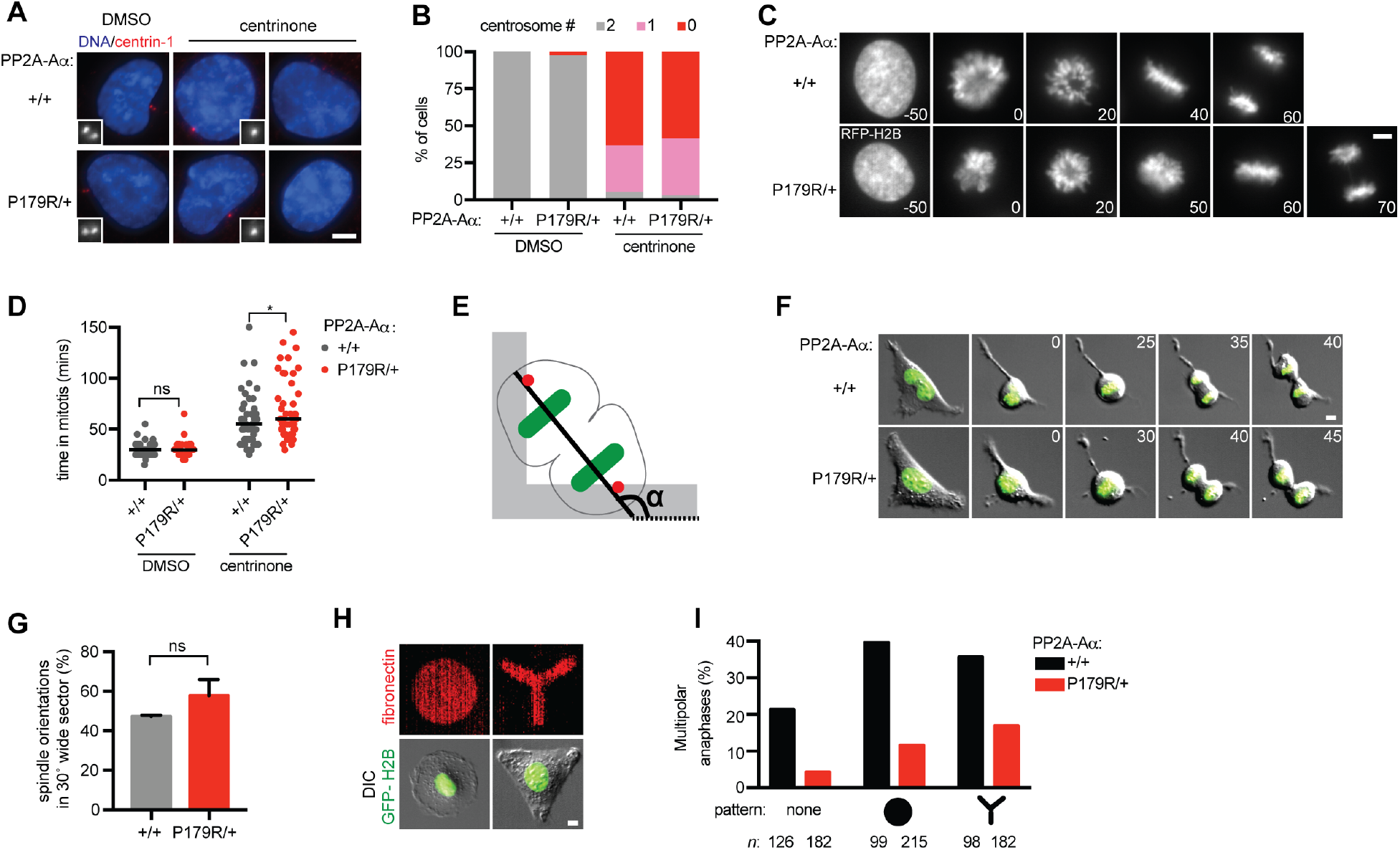
Cortical forces contribute to centrosome clustering efficiency in PP2A-Aα^P179R/+^ cells (A-B) Immunofluorescence analysis of WT (+/+) and PP2A-Aα^P179R/+^ (P179R/+) cells treated with centrinone. **(A)** Maximum intensity projections to score centrosome number. Inset, 2.5x enlargement. **(B)** Percentage of cells with the indicated number of centrosomes is plotted. **(C-D)** Cells were imaged live after DMSO or centrinone treatment. **(C)** RFP-H2B montage of centrinone treated cells. **(D)** Mitotic duration is plotted. Line, median; circle, cell. **(E)** Schematic of cell division on an L-shaped fibronectin pattern (grey). The angle (α) of chromosome segregation axis (solid line) relative to the reference X-axis (dashed line) is measured. Chromatin, green; Centrosome, red. **(F)** GFP-H2B and DIC merge time-lapse images. **(G)** Percentage of cells dividing within 30 degrees of the median angle is plotted. Mean + s.d. of two experiments. **(H-I)** Plk4 over-expressing cells were treated with dox and imaged on O- and Y-shaped fibronectin micro-patterns. **(H)** Representative images of interphase cells. **(I)** Incidence of multipolar anaphases. *n*, total cells analyzed. Scale bars, 5 μm. (C, F) Time (min) relative to nuclear envelope breakdown is indicated. (B, D) Representative quantifications were performed in triplicate. *, P <0.05, ns, not significant (P >0.05) Student’s *t*-test.

Centrosome clustering is a multi-step process (Rhys et al., 2018). In the first step, contacts between the actin network at the cell cortex and astral microtubules limit centrosome movement (Théry et al., 2005). To examine the cortical forces acting on centrosomes we plated cells on L-shaped fibronectin micro-patterns. Cortical forces, emanating from retraction fibers at the tips of the “L” shape, position the cell division axis along the hypotenuse of the ‘L’ (Fig. 5E-F). In 47% of WT cells, the orientation of the cell division axis was within 30 degrees of the hypotenuse. PP2A-Aα^P179R/+^ cells exhibited a similar proficiency to align the cell division axis (Fig. 5G) suggesting that centrosome positioning by cortical forces is similar in WT and PP2A-Aα^P179R/+^ cells.

To compare the impact of cortical forces in cells with centrosome amplification, we induced centrosome amplification by Plk4 overexpression and plated cells onto O- or Y-shaped fibronectin micro-patterns (Fig. 5H). Both shapes bias cells with supernumerary centrosomes towards multipolar cell divisions (Kwon et al., 2008). Indeed, in WT cells these patterns resulted in a 61% and 68% increase in the frequency of multipolar cell divisions on O- and Y-shaped patterns respectively (Fig. 5I). Interestingly, in PP2A-Aα^P179R/+^ cells the frequency of multipolar cell divisions was increased by 144% and 277% on O- and Y-shaped patterns respectively (Fig. 5I). Thus, centrosome clustering efficiency in PP2A-Aα^P179R/+^ cells is more sensitive to interphase cell shape. However, the frequency of multipolar cell divisions in PP2A-Aα^P179R/+^ cells was still reduced compared to WT cells (Fig. 5I). This result indicates that while cortical forces contribute to enhanced centrosome clustering in PP2A-Aα^P179R/+^ cells, the phenotype likely also arises from mechanisms unrelated to cell shape.

## Discussion

Here we have characterized a cancer-associated hotspot P179R mutation in PP2A-Aα that allows cells to more efficiently cluster supernumerary centrosomes. This finding is important because cells with supernumerary centrosomes only survive mitosis if they cluster centrosomes (Ganem et al., 2009). Our data suggests that a potential explanation for enrichment of PP2A-Aα mutations in tumors that experience WGD is to allow cells to survive the mitotic stress associated with centrosome amplification. PP2A-dependent dephosphorylation has myriad roles in cell proliferation, and our work points to a change in phospho-signaling that enables more robust centrosome clustering. Future work will be required to determine if there are additional impacts of hotspot mutations in the context of WGD, such as a compromised phospho-regulation of the G1/S checkpoint, which would allow cells to more readily proliferate after WGD.

Our work underscores the importance of considering PP2A-Aα mutations in a context that closely resembles the genetic change in tumors. Our data suggest that the enhanced centrosome clustering efficiency arises from haploinsufficiency in PP2A-Aα function, consistent with the known tumor suppressor role of PP2A (Pallas et al., 1990; Suganuma et al., 1988). If the primary impact of the mutation is to reduce PP2A-Aα functionality, then it is perhaps unexpected that hotspot missense mutations dominate the *PPP2R1A* mutational landscape rather than truncations. Several functions of PP2A-Aα remain in the P179R mutant, including association with the catalytic subunit and STRN regulatory subunits. Thus, the mutations may selectively impact the subset of PP2A holoenzymes with tumor suppressor functions.

Our work reveals that decreasing PP2A functionality increases centrosome clustering, opening new avenues of exploration to understand how cancer cells efficiently cluster centrosomes. Given that the only known function of PP2A is to dephosphorylate serine/threonine residues, our work indicates that fine-tuning phospho-regulation, in the form of a heterozygous PP2A-Aα mutation, allows a human cell to become more proficient at centrosome clustering without compromising mitotic fidelity. Because the mutation only partially reduces PP2A-Aα function, and a WT allele remains present in the cell, we predict that the observed phenotype is due to small changes in phosphorylation of substrates that control centrosome clustering. Future work will focus on identifying these changes, including phosphorylation sites that impact centrosome positioning and engagement during mitosis. Lastly, pharmacologic activators of PP2A have recently been described, and it will be important to investigate their anti-tumor efficacies in human cancers with specific oncogenic PP2A mutations (Ramaswamy et al., 2015).

## Acknowledgements

We thank Veronica Rodriguez-Bravo and Sun Joo Lee for experimental assistance, Prasad Jallepalli for helpful discussions and insightful critiques and Bryan Tsou for advice on Plk4 over-expression experiments. This work was supported by the Functional Genomics Initiative at Memorial Sloan Kettering (E.A.F. and A.K.), NIH/NIGMS GM125996 (E.A.F) and NCI CA214812 to A.K.

## Author contributions

Formal analysis and investigation: N.V.A., M.M.O., and P.C.; Conceptualization, experiment design and writing: N.V.A. and E.A.F.; Funding acquisition: E.A.F. and A. K.

## Materials and Methods

### Cell culture

hTERT WT cells (a gift from Alexey Khodjakov) and WT derivative cell lines were grown at 37°C in a humidified atmosphere with 5% CO_2_ in DMEM:F12 growth medium. Human embryonic kidney 293-T cells (ATCC) and Human embryonic kidney 293 cells (ATCC) were grown in DMEM growth medium. All media were supplemented with 10% fetal bovine serum (VWR Life Science), penicillin-streptomycin (Gemini Bio-Products), non-essential amino acids (Life Technologies), and L-glutamine (Sigma-Aldrich). Cell lines were authenticated upon initial receipt and upon initial generation by Short Tandem Repeat analysis at the Integrated Genomics Operations at Memorial Sloan Kettering Cancer Center. Cell lines were also tested for mycoplasma contamination by the Antibody and Bioresource core facility at Memorial Sloan Kettering Cancer Center.

### Cell line generation

To generate the PP2A-Aα^P179R/+^ WT cell lines, we used adeno-associated virus-mediated gene targeting. Q5 High-Fidelity DNA polymerase (New England Biolabs) was used to amplify two homology arms spanning exons 5 and 6 of *PPP2R1A* and cloned into a bacterial artificial chromosome (Life Technologies). The homology arms were cloned into vector pNX (Papi et al., 2005) flanking a *loxP*-Neo^R^-*loxP* cassette. Site-directed mutagenesis was used to introduce a C to G mutation at nucleotide 23280 in exon 5 of *PPP2R1A*. The *PPP2R1A* homology fragment was then cloned into pAAV (Berdougo et al., 2009) by NotI restriction enzyme digestion. Isolation of AAV particles, selection of stable transductants, and PCR screening were carried out as previously described (Berdougo et al., 2009). We determined the *PPP2R1A* genotype in isolated clones by genomic PCR amplification of *PPP2R1A* followed by Sanger sequencing. We isolated two clones (termed a and b), each with a heterozygous *PPP2R1A* P179R mutation. For each clone, the neomycin cassette was excised by infecting cells with adenovirus expressing Cre recombinase (Vector Development Laboratory, Baylor College of Medicine) at a multiplicity of infection of 80 plaque-forming units/cell. We identified single clones that were negative for the neomycin cassette but positive for the remnant loxP site by PCR amplification of endogenous *PPP2R1A*. Clone a was used for all experiments and, where indicated, clone b was also examined.

To generate *Tp53* knockout WT cell lines, codon-optimized Cas9 (Addgene 41815) and a guide RNA targeting *Tp53* (Wang et al., 2015) were transfected into cells using a Nucleofector 2b Device (Lonza) with the setting T023. Colonies were screened for p53 loss by western blot using an antibody for the N-terminus of p53 (Santa Cruz Biotechnology SC-126, clone DO-1). To generate cell lines expressing tetracycline-inducible Plk4, we used the lentiviral pLVX-Tight-Puro system (Clontech). Lentiviruses were generated in HEK 293T cells transfected with psPAX2 (Addgene plasmid 12260), pCMV-VSV-G (Addgene plasmid 8454) and either pLVX-Tet-On-Advanced (Clontech) or Plk4 cloned into pLVX-tight-puro (Wang et al., 2011) using Lipofectamine 2000 (Thermo Fisher Scientific) according to manufacturer’s instructions. 48 h after transfection, the virus containing suspension from HEK293T cells was filtered with a 0.45 μM filter (EMD Millipore), supplemented with polybrene (4 μg/mL, Sigma) and applied to Tp53-knockout WT cells or PP2A-Aα^P179R/+^ cells. Cells were first transduced with Plk4 in pLVX-tight-puro lentivirus and selected with puromycin (15 μg/mL, Thermo Fisher Scientific) for 1 week. Puromycin-resistant cells were then grown in Tetracycline-free serum (Clontech) and transduced with pLVX-Tet-On-Advanced lentivirus followed by G418 selection (500 μg/mL, Thermo Fisher Scientific) for 1 week. For each PP2A-Aα genotype, we chose a clone that yielded centrosome amplification in 60% of cells after 48 h doxycycline treatment (1 μg/mL).

We generated cells with fluorescent H2B by lentiviral transduction. H2B-RFP (Addgene plasmid 26001) or H2B-GFP (Addgene plasmid 25999) plasmids were co-transfected with psPAX2 and pCMV-VSV-G into HEK 293T cells. Supernatants were filtered, mixed with polybrene (4 μg/mL) and applied to target cells. We generated cells expressing GFP-PP2Aα by retroviral transduction as in (Foley et al., 2011) using WT PP2A-Aα (Foley et al., 2011) or P179R and R183W mutants, which were cloned via site-directed mutagenesis.

### Inhibitor treatments

Commercially available small molecules and chemical inhibitors were used as follows - cytochalasin D (0.2 μM, Santa Cruz Biotechnology), cytochalasin B (4 μM, Santa Cruz Biotechnology), blebbistatin (100 μM, Millipore Sigma), SB203580 (10 μM, Selleckchem), doxycycline (1 μg/mL, Fisher Scientific), nocodazole (3.3 μM, Santa Cruz Biotechnology), centrinone (200 nM, Tocris Bioscience), and Okadaic acid (200 nM, Santa Cruz Biotechnology). To induce whole genome doubling, cells were plated in a glass bottom dish (Cellvis) or on coverslips (Fisher) and, 24 h later, treated with cytochalasin D, cytochalasin B, or blebbistatin for 20 h. To wash out the inhibitor, the media was replaced six times over 1 h. *Tp53^+/+^* cells were released into SB203580, and imaged 20 h later. This inhibitor was omitted in *Tp53^−/−^* cells. Cells were then fixed and processed for immunofluorescence microscopy or imaged live by DIC and fluorescence microscopy. Plk4 induction with doxycycline was performed for 48 h. Unless otherwise indicated, ≥50 cells were analyzed per condition, per experiment and at least three experiments were analyzed per treatment condition. Where indicated, mitotic cell enrichment was performed using a 14 h treatment with nocodazole followed by manual detachment from the tissue culture dish.

### Live-cell imaging

Live imaging was performed on a Nikon Eclipse TiE inverted microscope equipped with a PI Piezo stage controller and Nikon Perfect Focus. Cells were maintained at 37 °C and 5% CO_2_ with a Stage Top Incubator with Flow Control and a Sub Stage Environmental Enclosure (Tokai Hit). Cells were imaged by DIC and fluorescence microscopy (GFP-Ex470 nm; Em522 nm or DsRed-Ex508 nm; Em620 nm). Images were acquired with a 20X or 40X objective and recorded on a deep-cooled, ultra-low noise sCMOS camera (Andor) at 16-bit depth using Elements software (Nikon). Image cropping was performed in Fiji (Schindelin et al., 2012).

### Immunofluorescence microscopy

For immunofluorescence microscopy of α-tubulin, C-Nap 1, and centrin-1, cells on coverslips were fixed in methanol at −20 °C for 10 min. For PP2A-Aα and B56α immunofluorescence, cells on coverslips were pre-extracted for 40 s at 37 °C in PEM buffer (100 mM K-PIPES pH 6.9, 10 mM EGTA, 1 mM MgCl_2_) with 0.5% Triton X-100 and 4 M glycerol. Coverslips were then fixed in PEM buffer with 3.7% formaldehyde and 0.2% Triton X-100 for 5 min at 37 °C. Coverslips were washed in TBS + 0.1% Triton X-100, blocked in same buffer with 2% donkey serum (Jackson Immunoresearch Laboratories) for 30 min, and stained with primary antibody for 3 h at room temperature. Species-specific secondary antibodies conjugated to Alexa Fluor 488, Rhodamine, Alexa Fluor 647 (Jackson Immunoresearch Laboratories), or IgG2a Alexa Fluor 594 (Thermo Fisher Scientific) were applied for 30 min at 0.6 μg/mL. Coverslips were mounted onto slides with Prolong Gold with DAPI (Thermo Fisher Scientific) and sealed. Imaging was performed on a DeltaVision Elite microscope running softWoRx software (GE Life Sciences). Images were acquired at 60X or 100X magnification with a PCO Edge CMOS Camera. Z-stacks were acquired with 0.2 μm spacing. Maximum intensity projection images were created in softWoRx. Pseudo coloring and cropping was performed in Fiji (Schindelin et al., 2012) and images were assembled in Illustrator (Adobe, San Jose, CA). Kinetochore/centromere intensity was calculated in Fiji using published analysis macros (Nijenhuis et al., 2014). 20 cells were analyzed per condition per experiment.

### Antibodies

Primary antibodies were used at 1 μg/mL for immunofluorescence incubations and 0.1 μg/mL for western blotting incubations. 2-4 μg of IgG was used for immunoprecipitation. Custom PP2A-Aα antibody was generated by immunizing rabbits with the peptide sequence of MAAADGDDSLY conjugated to matriculture Keyhole Limpet Hemocyanin (Thermo Scientific). Antigens were injected into New Zealand White rabbits using an institutionally approved protocol and animal-care facility (Pocono Rabbit Farm and Laboratory, Canadensis, PA). To purify peptide antibodies, antisera were diluted with 0.1 vol 10X PBS and incubated overnight with immunizing peptide covalently coupled to sulfo-link resin (Thermo Fisher Scientific). After washing with PBS, antibodies were eluted with 0.2 M glycine, pH 2.5 and subsequently neutralized with Tris pH 8.0, followed by dialysis into PBS. Custom GFP antibodies were raised against bacterially expressed GST-GFP. The serum was loaded onto a HiTrap NHS-activated HP column (GE Life Sciences) coupled to GST-GFP. Commercial and published antibodies used in this study include: B56α (immunofluorescence: BD Biosciences, 610615; western (Lee et al., 2017)), B56γ (Bethyl Laboratories, A300-814A), B56δ (Bethyl Laboratories, A301-100A), B56ε (Lee et al., 2017), B55α (Santa Cruz Biotechnology, SC-81606), PP2A-Aα/β (Santa Cruz Biotechnology, SC-6112), PP2A-C (BD Biosciences, 610555), centrin-1 (EMD Millipore, 04-1624), CREST Serum (Immunovision, HCT-0100; used at 1:5,000 dilution), α-tubulin (mouse DM1α; Abcam, ab7291), α-tubulin FITC conjugated (Sigma F2168), β-actin (Santa Cruz Biotechnology, SC-47778), p53 (Santa Cruz Biotechnology SC-126, clone DO-1), and C-Nap 1(a gift from Bryan Tsou).

### Mutation analysis in *PPP2R1A*

*PPP2R1A* mutations were quantified from publicly available studies on cBioPortal (Cerami et al., 2012; Gao et al., 2013) as of May 2018.

### Cell lysis, immunoprecipitation and phosphatase assays

Frozen cell pellets were suspended in buffer B (30 mM HEPES pH 7.8, 140 mM NaCl, 6 mM MgCl_2_, 5% glycerol) supplemented with 2 mM DTT, ProBlock Gold Mammalian Protease Inhibitor Cocktail (GoldBio Technology) and PhosSTOP (Roche) and maintained at 4 °C. Samples were lysed by nitrogen cavitation (Parr Instruments) for 5 min at 2,000 psi, and then centrifuged at 20,000 x *g* for 15 min. Proteins were immunoprecipitated with antibody bound to Protein A Dynabeads (ThermoFisher Scientific) for 1 h, washed three times with buffer B, and analyzed by western blot via chemiluminescence using an ImageQuant LAS500 (GE Life Sciences). Image cropping and intensity measurements were performed in Fiji (Schindelin et al., 2012) and images were assembled in Illustrator (Adobe, San Jose, CA). For phosphatase assays, cell pellets were suspended in buffer B with protease inhibitor and DTT. Lysates were mixed with protein G sepharose beads (GE Healthcare) bound to α-PP2A-Aα or control IgG. After washing in buffer B, samples were split and incubated with 200 nM Okadaic acid or DMSO. A portion was reserved for western blot analysis. The remainder was used in the Ser/Thr phosphatase assay (Promega). Final assay conditions were: 80 μM phosphopeptide, 50 mM imidazole pH 7.2, 0.2mM EGTA, 0.02% β-mercaptoethanol, 0.1 mg/mL BSA + 144 nM okadaic acid. Phosphatase activity was quantified by absorbance measured at 630 nm on a spectrophotometer.

### PP2A-Aα immunoprecipitation for SILAC mass spectrometry analysis

Wild type RPE-1 cells expressing GFP-PP2A-Aα (WT, P179R, or R183W) were grown in DMEM:F-12 (1:1) media for SILAC (ThermoFisher Scientific), supplemented with 10% dialyzed FBS (ThermoFisher Scientific), 1X penicillin-streptomycin (Gemini Bio-Products), 2.5 mM L-Glutamine (Sigma-Aldrich), 0.175 mM Arginine and 0.25 mM Lysine. Arg^0^ and Lys^0^ amino acids were purchased from Sigma. Arg^10^ (^13^C_6_/^15^N_4_ arginine) and Lys^8^ (^13^C_6_/^15^N_2_ lysine) amino acids were purchased from Cambridge Isotopes. After two weeks of passaging, incorporation of heavy amino acids was measured by mass spectrometry. Samples with 95% or greater incorporation of Lys^8^ and Arg^10^ were used in analyses. For immunoprecipitation, asynchronously growing cells were trypsinized, pelleted, and frozen. Cells were thawed and re-suspended in lysis buffer (180 mM NaCl, 50 mM sodium phosphate pH 7.4, 2 mM MgCl_2_, 0.1% Tween-20, 1X ProBlock Gold Mammalian Protease Inhibitor Cocktail (GoldBio Technology) and 1X Simple Stop 3 phosphatase inhibitor (GoldBio Technology)). All further steps were carried out at 4 °C. Cells were lysed by nitrogen cavitation (Parr Instruments) at 2,000 psi for 10 min, followed by centrifugation at 20,000 x *g* for 20 min. The supernatant was centrifuged once more. 5 mg protein was incubated with 125 μg α-GFP antibody covalently coupled to M270 magnetic resin (ThermoFisher Scientific) for 1 h with rotation. The resin was washed five times with lysis buffer. Bound proteins were eluted into 0.5 N NH_4_OH, 0.5 mM EDTA and dehydrated in a vacufuge (Eppendorf).

Dehydrated samples were suspended in 20 μL of NuPAGE LDS sample buffer (ThermoFisher Scientific). SILAC pairs were mixed, denatured at 90 °C for 2 min and separated on a 10% NuPAGE gel (ThermoFisher Scientific) at 120 V for 20 min. The gel was washed and silver stained (Pierce Silver Stain kit). Each lane was excised and destained using 30 mM potassium hexa-cyanoferrate (III) /100 mM sodium thiosulfate, washed and de-hydrated using a vacuum centrifuge. Proteins in the gel slabs were reduced, alkylated with iodoacetamide, dehydrated in a vacuum centrifuge. Gel slabs were then re-hydrated using 50 mM ammonium bicarbonate pH 8.4 (containing 0.04 μg sequencing grade modified porcine trypsin (Promega)) and incubated for 16 h at 37 °C. After proteolysis inhibition and tryptic peptide elution from the polyacrylamide slabs, eluates from each lane were pooled, lyophilized, and stored at −80 °C until analysis. The lyophilized peptide pellets were resuspended in 0.1% formic acid/ 3% acetonitrile and 5% of the solution was analyzed by LC/MS. The LC system consisted of a vented trap-elute setup (EasynLC1000, Thermo Fisher scientific) coupled to the Orbitrap Fusion mass spectrometer (Thermo Fisher Scientific, San Jose, CA) via a nano electrospray DPV-565 PicoView ion source (New Objective). The trap column was fabricated with a 5 cm × 150 μm internal diameter silica capillary with a 2 mm silicate frit, and pressure loaded with Poros R2-C18 10 μm particles (Life Technologies). The analytical column consisted of a 25 cm × 75 μm internal diameter column with an integrated electrospray emitter (New Objective), packed with ReproSil-Pur C18-AQ 1.9 μm particles (Dr. Maisch). Peptides were resolved over 90 min using a 3%–45% gradient acetonitrile/ 0.1% formic acid (buffer B) gradient in a water/0.1% formic acid (buffer A) at 250 nL/minute. Precursor ion scans were recorded from 400–2000 m/z in the Orbitrap (240,000 resolution at m/z 200) with an automatic gain control target set at 10^5^ ions and a maximum injection time of 50 ms. We used data-dependent mass spectral acquisition with monoisotopic precursor selection, ion charge selection (2–7), dynamic precursor exclusion (60 s, 20 ppm tolerance) and HCD fragmentation (normalized collision energy 35, isolation window 0.8 Th) using the top speed algorithm with a duty cycle of 2 s. Product ion spectra were recorded in the linear ion trap (“normal” scan rate, automatic gain control = 5000 ions, maximum injection time = 150 ms). Spectra were analyzed using MaxQuant Version1.5.2.8, searching within the human UniProt database (version 01/27-2016) with FDR <0.01. Intensity measurements were normalized to the PP2A-Aα intensity and then SILAC ratios were calculated. Four biological replicates were analyzed. P-values were calculated from a two-tailed student’s *t*-test for proteins identified in three or more experiments. Raw files are openly accessible via PRIDE accession number PXD010709.

---

**Figure S1.**
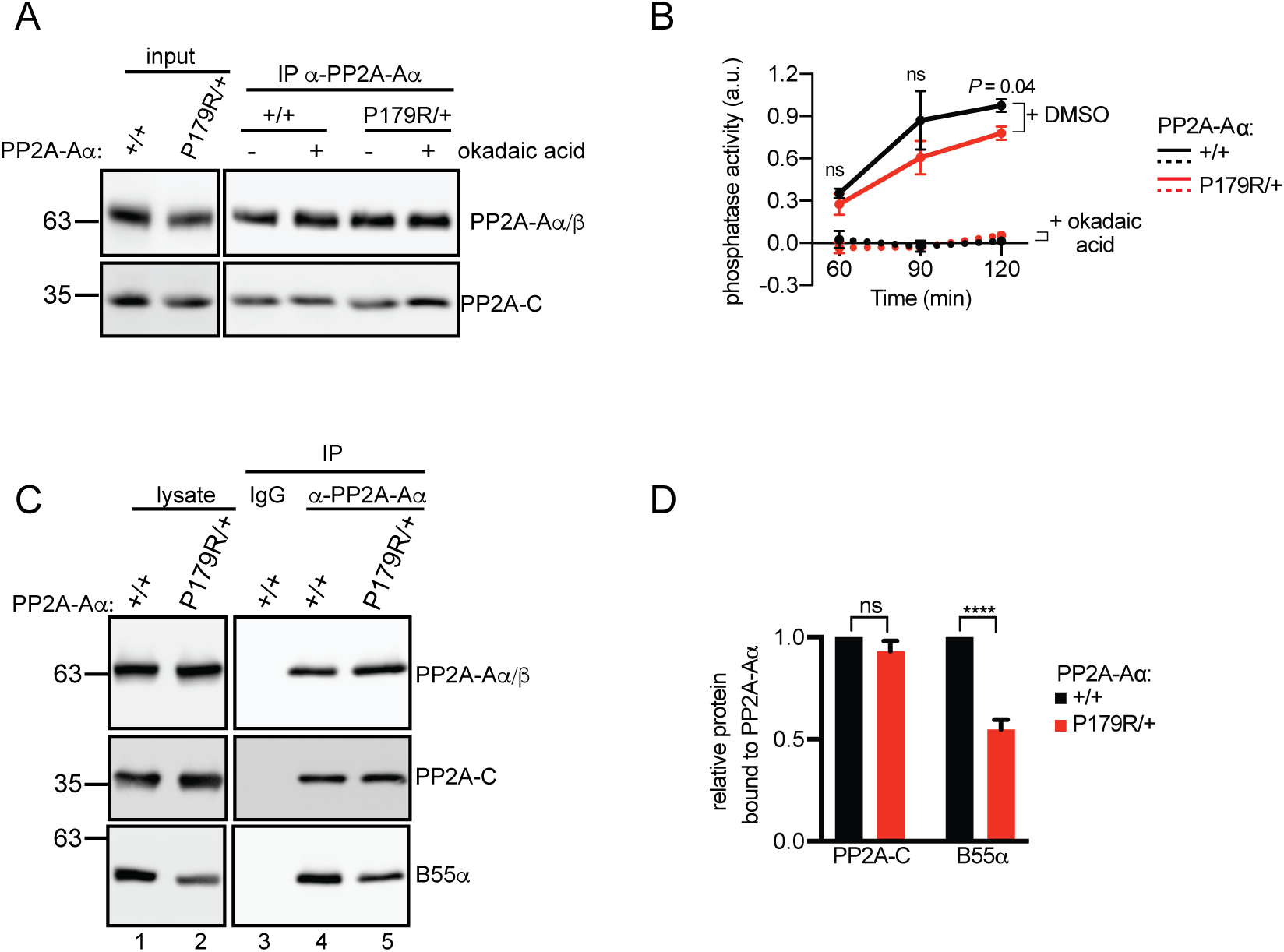
Analysis of PP2A activity and holoenzyme assembly in PP2A-Aα^P179R/+^ cells. **(A-B)** PP2A-Aα IPs in mitotic lysates from WT (+/+) or PP2A-Aα^P179R/+^ (P179R/+). Okadaic acid was included in wash steps as indicated. The reaction was split and (A) associated proteins were analyzed by western blot or (B) incubated with a phosphopeptide substrate. Phosphate release was quantified using a molybdate dye-based spectrophotometric assay. Absorbance values were normalized to the maximum value per experiment. Mean + s.d. from three experiments is plotted. **(C)** Western blot analysis of lysates (lanes 1-2) and IPs of control IgG (lane 3) or PP2A-Aα IgG (lane 4-5) in indicated cell lines **(D)** Plotted is the normalized mean ± s.e.m. of the experiment in (C) performed three times. ****, P <0.0005, ns, not significant (P >0.05) Student’s *t*-test.

**Figure S2.**
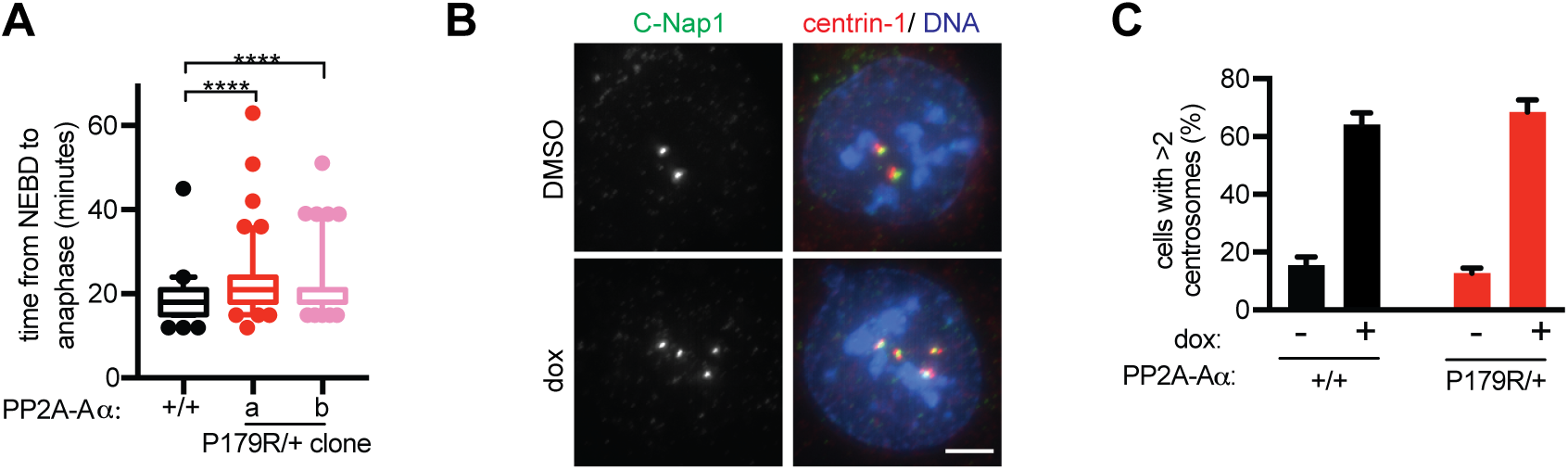
Analysis of mitotic duration in PP2A-Aα^P179R/+^ cells and centrosome amplification efficiency in Plk4-inducible cells. **(A)** WT (+/+) and PP2A-Aα^P179R/+^ (P179R/+) cells were imaged live and the time from nuclear envelope breakdown to anaphase onset was measured. Clones a and b are independently derived cell lines. A box-and-whisker plot is shown. Whiskers indicate the 5-95 percentile range and circles indicate cells outside of this range. The result is representative of two experiments. **(B-C)** *Tp53^−/−^* WT (+/+) and PP2A-Aα^P179R/+^ (P179R/+) cells with tet-inducible Plk4 expression were treated with dox or DMSO for 30 h, arrested in G2 with RO-3306 and dox or DMSO for 18 h, and processed for immunofluorescence. **(B)** Maximum intensity projections of C-Nap1 and a merge with centrin-1 and DNA. **(C)** Fraction of cells with >2 centrosomes (mean + s.d. from two experiments with 200 cells scored per condition, per experiment). Scale bar, 5 μm. a.u., arbitrary units; ****, P <0.0005, *, P <0.05, ns, not significant (P >0.05) Student’s *t*-test.

## References

Andreassen, P. R., Lohez, O. D., Lacroix, F. B. and Margolis, R. L. (2001). Tetraploid state induces p53-dependent arrest of nontransformed mammalian cells in G1. Mol. Biol. Cell 12, 1315–28.

Basto, R., Brunk, K., Vinadogrova, T., Peel, N., Franz, A., Khodjakov, A. and Raff, J. W. (2008). Centrosome amplification can initiate tumorigenesis in flies. Cell 133, 1032–42.

Berdougo, E., Terret, M.-E. and Jallepalli, P. V (2009). Functional dissection of mitotic regulators through gene targeting in human somatic cells. Methods Mol. Biol. 545, 21–37.

Bielski, C. M., Zehir, A., Penson, A. V., Donoghue, M. T. A., Chatila, W., Armenia, J., Chang, M. T., Schram, A. M., Jonsson, P., Bandlamudi, C., et al. (2018). Genome doubling shapes the evolution and prognosis of advanced cancers. Nat. Genet.

Carter, S. L., Cibulskis, K., Helman, E., McKenna, A., Shen, H., Zack, T., Laird, P. W., Onofrio, R. C., Winckler, W., Weir, B. A., et al. (2012). Absolute quantification of somatic DNA alterations in human cancer. Nat. Biotechnol. 30, 413–21.

Cerami, E., Gao, J., Dogrusoz, U., Gross, B. E., Sumer, S. O., Aksoy, B. A., Jacobsen, A., Byrne, C. J., Heuer, M. L., Larsson, E., et al. (2012). The cBio cancer genomics portal: an open platform for exploring multidimensional cancer genomics data. Cancer Discov. 2, 401–4.

Chan, J. Y. (2011). A clinical overview of centrosome amplification in human cancers. Int. J. Biol. Sci. 7, 1122–44.

Chen, W., Possemato, R., Campbell, K. T., Plattner, C. A., Pallas, D. C. and Hahn, W. C. (2004). Identification of specific PP2A complexes involved in human cell transformation. Cancer Cell 5, 127–36.

Cho, U. S. and Xu, W. (2007). Crystal structure of a protein phosphatase 2A heterotrimeric holoenzyme. Nature 445, 53–57.

Cooper, J. A. (1987). Effects of cytochalasin and phalloidin on actin. J. Cell Biol.

Cuenda, A., Rouse, J., Doza, Y. N., Meier, R., Cohen, P., Gallagher, T. F., Young, P. R. and Lee, J. C. (1995). SB 203580 is a specific inhibitor of a MAP kinase homologue which is stimulated by cellular stresses and interleukin-1. FEBS Lett. 364, 229–233.

Dewhurst, S. M., McGranahan, N., Burrell, R. A., Rowan, A. J., Grönroos, E., Endesfelder, D., Joshi, T., Mouradov, D., Gibbs, P., Ward, R. L., et al. (2014). Tolerance of whole-genome doubling propagates chromosomal instability and accelerates cancer genome evolution. Cancer Discov. 4, 175–85.

Drosopoulos, K., Tang, C., Chao, W. C. H. and Linardopoulos, S. (2014). APC/C is an essential regulator of centrosome clustering. Nat. Commun. 5,.

Foley, E. A., Maldonado, M. and Kapoor, T. M. (2011). Formation of stable attachments between kinetochores and microtubules depends on the B56-PP2A phosphatase. Nat Cell Biol 13, 1265–71.

Ganem, N. J., Godinho, S. A. and Pellman, D. (2009). A mechanism linking extra centrosomes to chromosomal instability. Nature 460, 278–282.

Gao, J., Aksoy, B. A., Dogrusoz, U., Dresdner, G., Gross, B., Sumer, S. O., Sun, Y., Jacobsen, A., Sinha, R., Larsson, E., et al. (2013). Integrative analysis of complex cancer genomics and clinical profiles using the cBioPortal. Sci. Signal. 6, pl1.

Haesen, D., Abbasi Asbagh, L., Derua, R., Hubert, A., Schrauwen, S., Hoorne, Y., Amant, F., Waelkens, E., Sablina, A. and Janssens, V. (2016). Recurrent PPP2R1A Mutations in Uterine Cancer Act through a Dominant-Negative Mechanism to Promote Malignant Cell Growth. Cancer Res. 76, 5719–5731.

Hertz, E. P. T., Kruse, T., Davey, N. E., López-Méndez, B., Sigurðsson, J. O., Montoya, G., Olsen, J. V and Nilsson, J. (2016). A Conserved Motif Provides Binding Specificity to the PP2A-B56 Phosphatase. Mol. Cell 63, 686–95.

Hyodo, T., Ito, S., Asano-Inami, E., Chen, D. and Senga, T. (2016). A regulatory subunit of protein phosphatase 2A, PPP2R5E, regulates the abundance of microtubule crosslinking factor 1. FEBS J. 283, 3662–3671.

Jeong, A. L., Han, S., Lee, S., Su Park, J., Lu, Y., Yu, S., Li, J., Chun, K.-H., Mills, G. B. and Yang, Y. (2016). Patient derived mutation W257G of PPP2R1A enhances cancer cell migration through SRC-JNK-c-Jun pathway. Sci. Rep. 6, 27391.

Kleylein-Sohn, J., Westendorf, J., Le Clech, M., Habedanck, R., Stierhof, Y.-D. and Nigg, E. A. (2007). Plk4-induced centriole biogenesis in human cells. Dev. Cell 13, 190–202.

Kwon, M., Godinho, S. A., Chandhok, N. S., Ganem, N. J., Azioune, A., Thery, M. and Pellman, D. (2008). Mechanisms to suppress multipolar divisions in cancer cells with extra centrosomes. Genes Dev. 22, 2189–203.

Kwon, M., Bagonis, M., Danuser, G. and Pellman, D. (2015). Direct Microtubule-Binding by Myosin-10 Orients Centrosomes toward Retraction Fibers and Subcortical Actin Clouds. Dev. Cell.

Leber, B., Maier, B., Fuchs, F., Chi, J., Riffel, P., Anderhub, S., Wagner, L., Ho, A. D., Salisbury, J. L., Boutros, M., et al. (2010). Proteins required for centrosome clustering in cancer cells. Sci. Transl. Med. 2, 33ra38.

Nijenhuis, W., Vallardi, G., Teixeira, A., Kops, G. J. P. L. P. L. and Saurin, A. T. (2014). Negative feedback at kinetochores underlies a responsive spindle checkpoint signal. Nat. Cell Biol. 16, 1257–64.

Pallas, D. C., Shahrik, L. K., Martin, B. L., Jaspers, S., Miller, T. B., Brautigan, D. L. and Roberts, T. M. (1990). Polyoma small and middle T antigens and SV40 small t antigen form stable complexes with protein phosphatase 2A. Cell 60, 167–176.

Papi, M., Berdougo, E., Randall, C. L., Ganguly, S. and Jallepalli, P. V (2005). Multiple roles for separase auto-cleavage during the G2/M transition. Nat. Cell Biol. 7, 1029–35.

Peel, N., Stevens, N. R., Basto, R. and Raff, J. W. (2007). Overexpressing Centriole-Replication Proteins In Vivo Induces Centriole Overduplication and De Novo Formation. Curr. Biol.

Quintyne, N. J., Reing, J. E., Hoffelder, D. R., Gollin, S. M. and Saunders, W. S. (2005). Spindle multipolarity is prevented by centrosomal clustering. Science 307, 127–9.

Ramaswamy, K., Spitzer, B. and Kentsis, A. (2015). Therapeutic Re-Activation of Protein Phosphatase 2A in Acute Myeloid Leukemia. Front. Oncol.

Rhys, A. D., Monteiro, P., Smith, C., Vaghela, M., Arnandis, T., Kato, T., Leitinger, B., Sahai, E., McAinsh, A., Charras, G., et al. (2018). Loss of E-cadherin provides tolerance to centrosome amplification in epithelial cancer cells. J. Cell Biol.

Ring, D., Hubble, R. and Kirschner, M. (1982). Mitosis in a cell with multiple centrioles. J. Cell Biol. 94, 549–56.

Schindelin, J., Arganda-Carreras, I., Frise, E., Kaynig, V., Longair, M., Pietzsch, T., Preibisch, S., Rueden, C., Saalfeld, S., Schmid, B., et al. (2012). Fiji: an open-source platform for biological-image analysis. Nat. Methods 9, 676–682.

Straight, A. F., Cheung, A., Limouze, J., Chen, I., Westwood, N. J., Sellers, J. R. and Mitchison, T. J. (2003). Dissecting temporal and spatial control of cytokinesis with a myosin II inhibitor. Science (80-.).

Suganuma, M., Fujiki, H., Suguri, H., Wakamatsu, K., Yamada, K., Sugimura, T., Yoshizawa, S., Nakayasu, M., Hirota, M. and Ojika, M. (1988). Okadaic acid: an additional non-phorbol-12-tetradecanoate-13-acetate-type tumor promoter. Proc. Natl. Acad. Sci.

Théry, M., Racine, V., Pépin, A., Piel, M., Chen, Y., Sibarita, J.-B. and Bornens, M. (2005). The extracellular matrix guides the orientation of the cell division axis. Nat. Cell Biol. 7, 947–953.

Wang, W. J., Soni, R. K., Uryu, K. and Tsou, M. F. B. (2011). The conversion of centrioles to centrosomes: Essential coupling of duplication with segregation. J. Cell Biol. 193, 727–739.

Wang, W. J., Acehan, D., Kao, C. H., Jane, W. N., Uryu, K. and Tsou, M. F. B. (2015). De novo centriole formation in human cells is error-prone and does not require SAS-6 self-assembly. Elife 4,.

Wlodarchak, N. and Xing, Y. (2016). PP2A as a master regulator of the cell cycle. Crit. Rev. Biochem. Mol. Biol.

Wong, Y. L., Anzola, J. V., Davis, R. L., Yoon, M., Motamedi, A., Kroll, A., Seo, C. P., Hsia, J. E., Kim, S. K., Mitchell, J. W., et al. (2015). Reversible centriole depletion with an inhibitor of Polo-like kinase 4. Science (80-.). 348, 1155–1160.

Xu, Y., Xing, Y., Chen, Y., Chao, Y., Lin, Z., Fan, E., Yu, J. W., Strack, S., Jeffrey, P. D. and Shi, Y. (2006). Structure of the protein phosphatase 2A holoenzyme. Cell 127, 1239–1251.

Xu, Y., Chen, Y., Zhang, P., Jeffrey, P. D. and Shi, Y. (2008). Structure of a protein phosphatase 2A holoenzyme: insights into B55-mediated Tau dephosphorylation. Mol Cell 31, 873–885.

Yang, Z., Lončarek, J., Khodjakov, A. and Rieder, C. L. (2008). Extra centrosomes and/or chromosomes prolong mitosis in human cells. Nat. Cell Biol. 10, 748–751.

Zack, T. I., Schumacher, S. E., Carter, S. L., Cherniack, A. D., Saksena, G., Tabak, B., Lawrence, M. S., Zhang, C.-Z., Wala, J., Mermel, C. H., et al. (2013). Pancancer patterns of somatic copy number alteration. Nat. Genet. 45, 1134–1140.

Zhou, J., Pham, H. T., Ruediger, R. and Walter, G. (2003). Characterization of the Aalpha and Abeta subunit isoforms of protein phosphatase 2A: differences in expression, subunit interaction, and evolution. Biochem. J. 369, 387–98.

